# A novel Toxoplasma IMC sutures-associated protein regulates suture protein targeting and colocalizes with membrane trafficking machinery

**DOI:** 10.1101/2021.06.22.449166

**Authors:** Jessica H. Chern, Andy S. Moon, Allan L. Chen, Jihui Sha, James A. Wohlschlegel, Peter J. Bradley

## Abstract

The cytoskeleton of *Toxoplasma gondii* is composed of the inner membrane complex (IMC) and an array of underlying microtubules that provide support at the periphery of the parasite. Specific subregions of the IMC carry out distinct roles in replication, motility, and host cell invasion. Building on our previous *in vivo* biotinylation (BioID) experiments of the IMC, we identify here a novel protein that localizes to discrete punctae that are embedded in the parasite’s cytoskeleton along the IMC sutures. Gene knockout analysis shows that loss of the protein results in defects in cytoskeletal suture protein targeting, cytoskeletal integrity, parasite morphology, and host cell invasion. We then use deletion analyses to identify a domain in the N-terminus of the protein that is critical for both localization and function. Finally, we use the protein as bait for *in vivo* biotinylation which identifies several other proteins that colocalize in similar spot-like patterns. These putative interactors include several proteins that are implicated in membrane trafficking and are also associated with the cytoskeleton. Together, this data reveals an unexpected link between the IMC sutures and membrane trafficking elements of the parasite and suggests that the sutures punctae are likely a portal for trafficking cargo across the IMC.

## Introduction

The phylum Apicomplexa is composed of obligate intracellular parasites that cause substantial disease in humans and animals worldwide (1). The most prominent apicomplexans that infect humans are *Toxoplasma gondii*, which causes disease in immunocompromised individuals and neonates, *Plasmodium spp*., the causative agents of malaria, and *Cryptosporidium spp*., which causes diarrheal diseases in children (2–4). Important animal pathogens include *Neospora caninum*, which causes abortion in cattle and neurological disease in dogs, and *Eimeria spp*. which causes disease in poultry (5). These parasites share a number of unique organelles that enable them to infect and replicate within their mammalian host cells (6). Because these organelles and many of their constituents are unique to the pathogens, they make attractive targets for the development of therapeutics that can specifically target the parasite.

One of these organelles is the inner membrane complex (IMC), which lies beneath the parasite’s plasma membrane and consists of both membrane and cytoskeletal elements (7). The IMC is additionally supported by a series of microtubules that are tethered to the basket shaped conoid at the apical end of the parasite and extend nearly the length of the cell. The IMC is known to carry out three major functions in infection of host cells and intracellular replication. First, it hosts the glideosome, an actin-myosin motor that interacts with adhesins secreted onto the parasite’s surface for motility and invasion (8). Second, it serves as a scaffold for the formation of daughter cells via the internal budding process known as endodyogeny (6). Finally, the apical cap portion of the organelle has recently been shown to support the conoid, a microtubule-based structure at the extreme apex of the cell which controls the release of secretory proteins for host cell invasion (9, 10). While these important activities of the IMC have been described and some of the key players identified, the precise roles of many of the constituents of the IMC have yet to be determined.

The IMC is able to carry out its diverse functions by partitioning the organelle into distinct subcompartments, each containing their own cargo of proteins (6, 7). The glideosome components that power motility are localized to the membrane vesicles of the IMC body and apical cap, with the motor facing the plasma membrane to tether to secreted micronemal adhesins (8). The apical cap portion of the organelle hosts the AC9/AC10/ERK7 complex, which regulates the stability of the conoid that is essential for release of the micronemes and rhoptries for attachment and penetration, respectively (9, 10). During endodyogeny, distinct groups of IMC proteins are synthesized in a “just in time” approach in which proteins are sequentially synthesized and added onto the membranes and cytoskeleton of forming daughter buds for the replication of new cells (11). The membrane vesicles of the IMC body are arranged into rectangular plates that are stitched together by the IMC suture proteins, which appear to be important for maintaining parasite shape and ensuring faithful replication (12, 13). IMC suture proteins are also associated with the parasite’s cytoskeletal network, although how they are arranged into their rectangular pattern and are tethered to the cytoskeleton is unknown. The basal complex is at the extreme base of the IMC and plays key roles in the expansion of the forming daughter buds and constriction of the daughters during the final stages of division (14). Lastly, the apical annuli are a series of five discrete spots anchored to the cytoskeleton between the apical cap and the parasite body, which may serve as pores for the transfer of nutrients or the removal of waste across the IMC (15).

One important advance in the discovery of many of the IMC proteins is the use of *in vivo* biotinylation (BioID) with bait proteins targeted to the various subcompartments, which can be used to identify new proximal and interacting proteins labeled in each location (12, 15–17). In this study, we characterize a protein identified in our previous IMC BioID experiments that surprisingly localizes to a series of punctae that colocalize with the IMC sutures and associate with the cytoskeleton (12, 13). Gene knockout of the protein shows that it is important for trafficking of a subgroup of the cytoskeletal suture proteins and integrity of the cytoskeleton, which results in defects in parasite morphology, replication, and host cell invasion. Deletion analyses demonstrate that the coiled-coil domains of the protein are surprisingly dispensable, but a conserved N-terminal region is essential for localization and function. We then use the protein as bait for BioID experiments which identifies several other proteins that colocalize with these punctae and are associated with the cytoskeleton. Among these are proteins that are implicated in membrane transport, suggesting that the IMC sutures punctae tether vesicle trafficking machinery for delivery of cargo into or across the IMC.

## Results

### TgGT1_202220 localizes to distinct cytoskeletal punctae along the IMC sutures

Our previous BioID experiments using IMC proteins as bait identified a large number of candidate IMC proteins, many of which have been verified by epitope tagging (9, 12, 13). One hit that was identified in BioID experiments using both AC9 and ISP3 proteins as bait was TgGT1_202220. TgGT1_202220 has a predicted mass of 122kDa and lacks identifiable protein domains other than two internal coiled-coil (CC) domains (Fig. 1A) (18). TgGT1_202220 is restricted to *T. gondii* and its closest relatives, as orthologues are present in *H. hammondi, N. caninum*, and *B. besnoiti*, but not in *Eimeria spp* or *Sarcocystis spp* (Fig. S1) (19). The highest regions of similarity appeared to be within the CC domains as well as in the N and C-terminal regions of the protein outside of the CC domains. To determine the localization of TgGT1_202220, we endogenously tagged the protein with a 3xHA tag which showed a series of faint spots in the parasite’s cytoplasm, some of which were close to background staining (Fig. 1B). In parasites that were dividing by endodyogeny, the spots appeared brighter in agreement with its cell cycle expression data (Fig. 1C) (11). To better assess localization of the protein, we used a spaghetti monster HA epitope tag (smHA) which enhances detection via its ten HA tags buried in a non-fluorescent GFP backbone (20). This revealed a substantially enhanced signal of the cytoplasmic spots (Fig. 1D), which again were brighter in dividing parasites (Fig. 1E).

**Figure 1.**
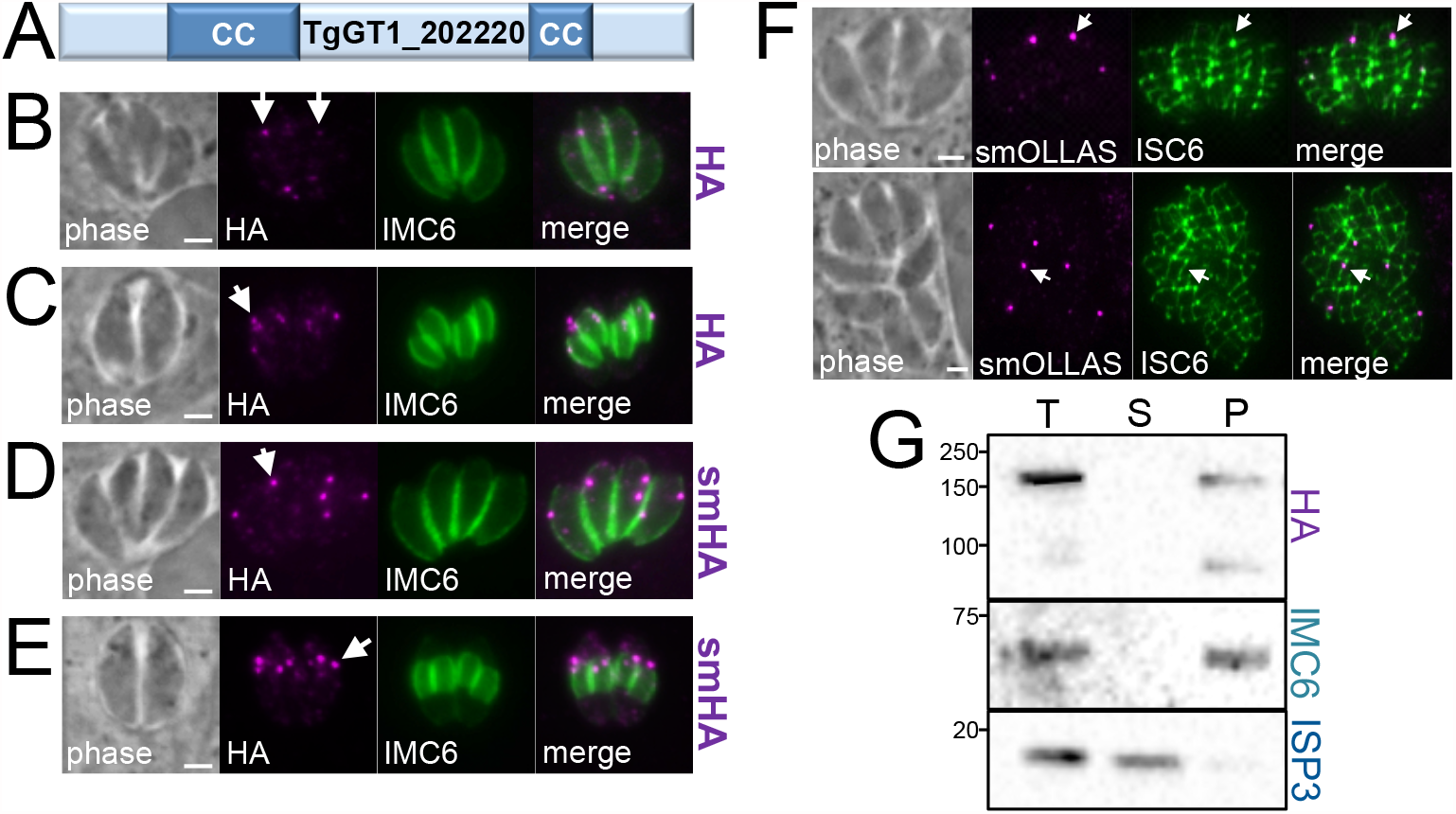
TgGT1_202220 localizes to cytoplasmic spots along the IMC sutures and is associated with the cytoskeleton. A) Diagram of TgGT1_202220 showing two predicted coiled-coil (CC) domains. B) IFA of endogenously 3xHA-tagged TgGT1_202220 showing several faint punctae in the cytoplasm (arrows). Magenta: mouse anti-HA; Green: rabbit anti-IMC6. C) IFA showing that the cytoplasmic punctae appear near the apical ends of the developing daughter buds during endodyogeny (arrow). Magenta: mouse anti-HA; Green: rabbit anti-IMC6. D) IFA showing endogenous tagging of TgGT1_202220 with spaghetti-monster HA (smHA) enables better detection of the cytoplasmic punctae (arrow). Magenta: mouse anti-HA; Green: rabbit anti-IMC6. E) TgGT1_202220 tagged with smHA also shows enhanced staining near the developing daughter buds (arrow). Magenta: mouse anti-HA; Green: rabbit anti-IMC6. F) IFA showing that smOLLAS-tagged TgGT1_202220 punctae colocalize with the IMC sutures (ISC6-3xHA). The arrow in the top panel shows a punctae on a longitudinal suture. The arrow in the bottom panel points to a punctae on a transverse suture. Magenta: rat anti-OLLAS; Green: mouse anti-HA. G) Western blot analysis of TX-100 detergent fractionation shows that TgGT1_202220 partitions to the cytoskeletal pellet with the alveolin IMC6 and is not released like the membrane-associated IMC protein ISP3. All scale bars in the figure are 2 µm.

Since the spotted localization of TgGT1_202220 differs from the peripheral localization of most IMC proteins, we examined structures associated with the IMC including the cortical microtubules, apical annuli, and IMC sutures. While we did not observe colocalization of the punctae with the microtubules or apical annuli (Fig. S2), the spots did colocalize with the IMC sutures (Fig. 1F). The signal was most frequently detected on the longitudinal sutures just adjacent to the intersection of the longitudinal and transverse sutures, but was sometimes seen on the transverse sutures as well. We then used detergent fractionation to determine if the protein is tethered to the parasite’s cytoskeleton, using IMC6 and ISP3 as controls for the cytoskeletal and membrane fractions, respectively (Fig. 1G) (12). While the protein reproducibly suffered from some breakdown during fractionation, the signal clearly partitioned with the insoluble cytoskeleton and was not released by detergent extraction. Together, these results indicate that TgGT1_202220 associates with the cytoskeleton at the IMC sutures and we thus named the protein IMC sutures-associated protein 1 (ISAP1).

### Disruption of ISAP1 substantially affects the parasite’s lytic cycle

ISAP1 was assigned a fitness score of −3.49 in the *Toxoplasma* genome-wide CRISPR/Cas9 screen (21), suggesting that the protein is either important for fitness or essential. To directly assess ISAP1 function, we used CRISPR/Cas9 to disrupt its gene from the HA epitope-tagged strain (22). IFA analysis of a clonal isolate of the knockout showed a loss of the HA staining and the knockout was confirmed by PCR (Fig. 2A, B). We then complemented the Δ*isap1* parasites with the full length ISAP1 gene driven from the ISC6 promoter, which restored the spot-like pattern by IFA and displayed similar levels of expression as the wild-type tagged strain (Fig. 2C, D) (13). These results demonstrated that ISAP1 can be disrupted and that its staining pattern and protein levels can be restored by complementation.

**Figure 2.**
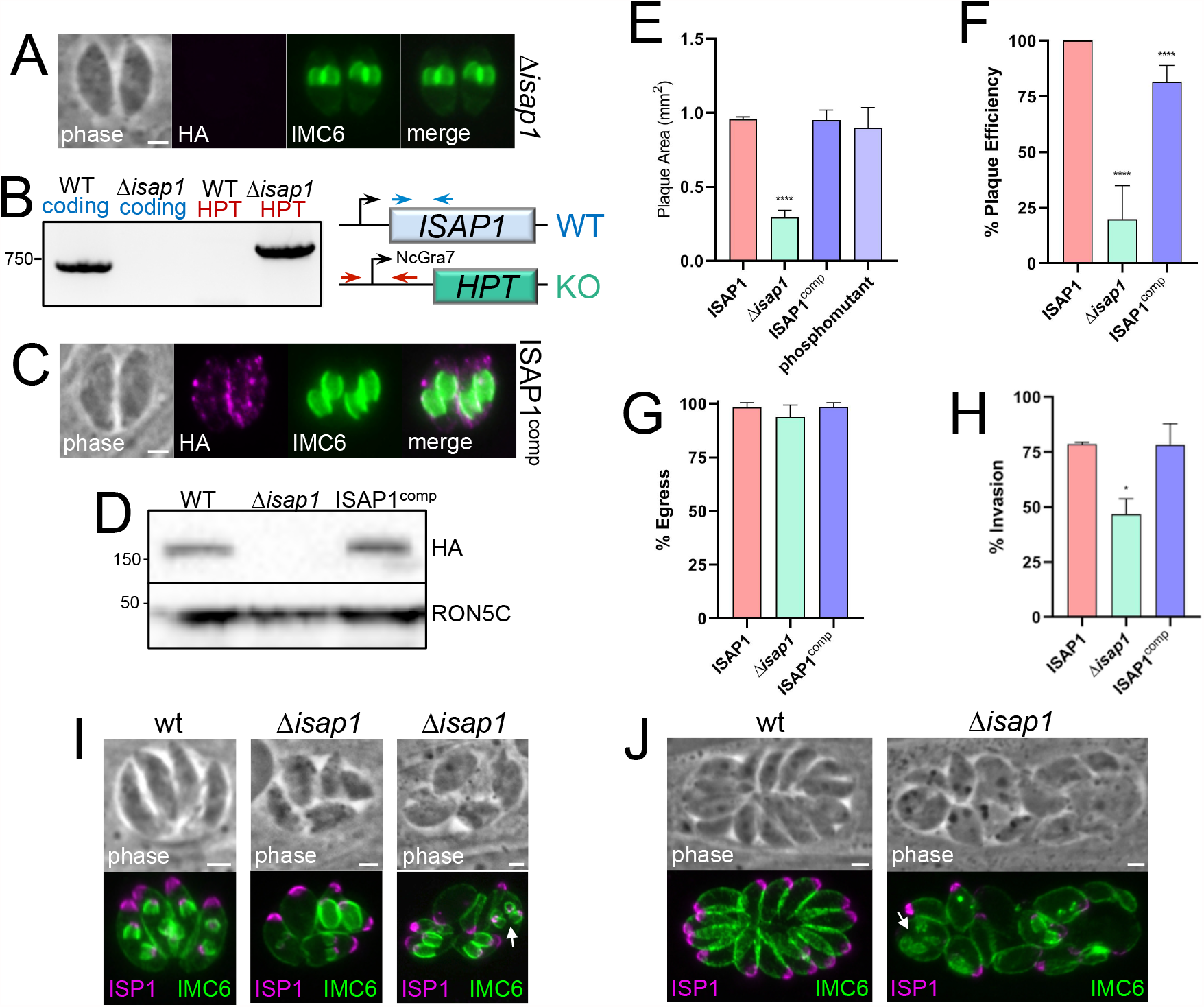
Gene knockout of ISAP1. A) IFA showing lack of HA staining in Δ*isap1* parasites. Magenta: mouse anti-HA; Green: rabbit anti-IMC6. Scale bar = 2 µm. B) PCR and diagram showing the Δ*isap1* strain contains the correct amplicon for the replacement of ISAP1 and lacks the ISAP1 coding amplicon. Primer positions are shown with arrows and amplicons agree with the anticipated sizes for the knockout using wild-type genomic DNA as a control. C) Complementation with the ISAP1 coding sequence driven from the ISC6 promotor restores the spot-like pattern similar to the wild-type protein. Magenta: mouse anti-HA; Green: rabbit anti-IMC6. Scale bar = 2 µm. D) Western blot analysis of HA-tagged, Δ*isap1* and complemented strains shows that complementation restores levels of the ISAP1 protein similar to that seen for the HA-tagged strain. Plaque assay showing disruption of ISAP1 results in a 69% decrease in plaque size (****, *P* < 0.0001). The defect is rescued by complementation (ISAP1^comp^). The ISAP1 phosphomutant (Fig. S3) also rescues the plaque defect. F) Graph showing an 80% reduction in plaque efficiency of Δ*isap1* parasites, which is mostly rescued upon complementation (****, *P* < 0.0001). G) Ionophore induced egress is not significantly affected in Δ*isap1* parasites. H) Host cell invasion is reduced by 42% in Δ*isap1* parasites (*\*, P* > 0.05). I) IFA at 24 hours post infection showing that loss of ISAP1 results in swollen parasites that have dysregulated endodyogeny and morphological defects. An arrow points to a parasite with four daughter buds. Magenta: mouse anti-ISP1; Green: rabbit anti-IMC6. Scale bar = 2 µm. J) IFA at 32 hours post infection showing more severe defects in morphology and daughter cell formation. Arrow points to four daughter buds in a maternal parasite. Magenta: mouse anti-ISP1; Green: rabbit anti-IMC6. Scale bar = 2 µm.

In agreement with its negative fitness score, the knockout parasites grew poorly, which we assessed via plaque assay. These experiments showed a 69% reduction in plaque size, which was fully rescued by complementation (Fig. 2E). To address phosphorylation of ISAP1, we additionally complemented the Δ*isap1* strain with a phosphomutant copy in which 7 phosphosites identified by phosphoproteomics were mutated to alanine. This phosphomutant localized to similar spots and fully rescued the plaque defect, demonstrating that these phosphosites are not important for function (Fig. 2E, Fig. S3). In addition to plaque size, we observed an 80% reduction in plaque efficiency, which was mostly rescued by complementation (Fig. 2F). The reductions in plaque size and efficiency indicated that some element of the lytic cycle is disrupted in the Δ*isap1* strain. To further characterize the defect, we individually examined host cell invasion, intracellular replication, and egress. While no defect was observed in egress, we did see a significant loss of invasive capability of the knockout (Fig. 2G, H). We next examined replication and noticed that the Δ*isap1* parasites often appeared swollen and many vacuoles had parasites with morphological defects, more than two daughter buds, or a loss of coordinated endodyogeny (Fig. 2I, J). We also examined an array of other organelles including the mitochondrion, apicoplast, rhoptries, and micronemes and only observed defects in those parasites with the most severe morphological problems, which suggested these were the result of parasite death (data not shown).

### Deletion of ISAP1 affects cytoskeletal integrity and targeting of a subset of the cytoskeletal suture proteins

In the course of examining the morphological defects of the Δ*isap1* parasites, we also noticed that many of the parasites had an apparent gap in the alveolin network proteins IMC1 and IMC6, suggesting a breach in the cytoskeleton (Fig. 3A). To further explore this and to begin to examine the IMC suture proteins, we tagged the cytoskeletal suture protein TSC2 in Δ*isap1* parasites, which targeted correctly to the sutures but also showed the apparent breach of the cytoskeleton (Fig. 3B). We also examined several other suture proteins including ISCs 1-6 and TSCs 3-6. While ISCs 3/5/6 and TSCs 3-6 appeared to traffic to the sutures correctly (Fig. 3C, D), ISC1 and ISC2 were mislocalized to a cytoplasmic spotted pattern and ISC4 was absent altogether (tagging was verified by PCR) (Fig. 3E-H). Together, this data indicates that ISAP1 is necessary for the correct targeting of ISCs 1/2/4 and for the integrity of the cytoskeletal meshwork of the IMC.

**Figure 3.**
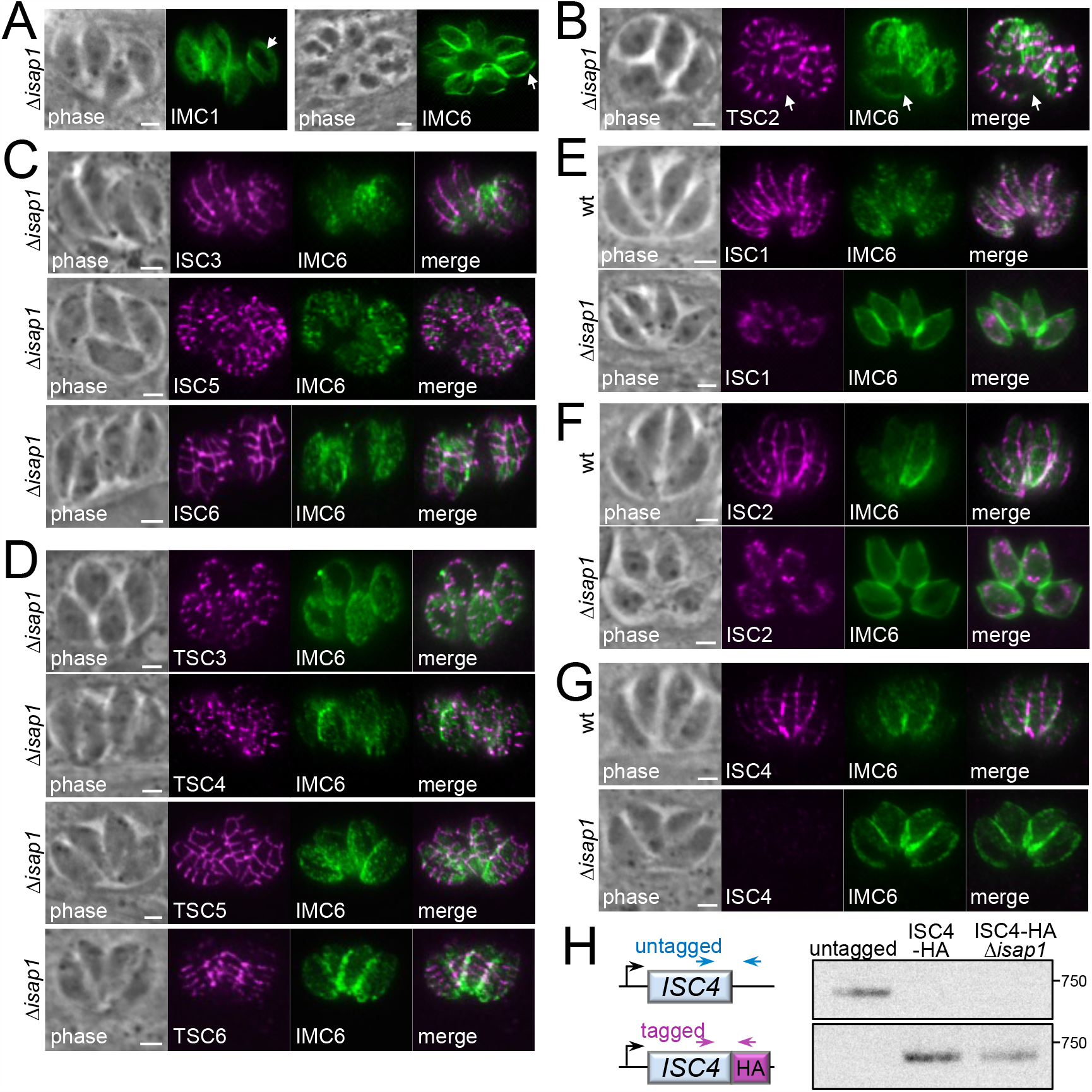
Loss of ISAP1 affects the integrity of the IMC cytoskeleton and results in mistargeting of a subgroup of cytoskeletal IMC suture proteins. A) IFA of IMC1 and IMC6 in Δ*isap1* parasites showing gaps in the IMC cytoskeleton (arrows). The more internal IMC1/IMC6 staining pattern throughout Fig. 3 is due to the planes imaged for visualizing the gaps or sutures at the periphery of the parasite. Green: mouse anti-IMC1 and rabbit anti-IMC6. B) IFA with the cytoskeletal transverse sutures protein TSC2 also shows the gaps in the Δ*isap1* cytoskeleton (arrows). TSC2 targets correctly to the transverse sutures. Magenta: mouse anti-HA; Green: rabbit anti-IMC6. C) IFA of Δ*isap1* parasites showing that HA-tagged ISC3, ISC5 and ISC6 target correctly to the sutures. Magenta: mouse anti-HA; Green: rabbit anti-IMC6. D) IFA of Δ*isap1* parasites showing that HA-tagged TSCs 3/4, Ty-tagged TSC5, and Myc-tagged TSC6 also target correctly to the sutures. Magenta: mouse anti-HA, mouse anti-Ty, mouse anti-Myc; Green: rabbit anti-IMC6. E-G) IFA showing that ISC1 and ISC2 are mislocalized to cytoplasmic spots and that ISC4 is absent in Δ*isap1* parasites. Magenta: rat anti-ISC1, rat anti-ISC2, mouse anti-HA (ISC4); Green: rabbit anti-IMC6. H) Strategy and agarose gel analysis showing PCR verification of correct tagging of ISC4 in Δ*isap1* parasites. All scale bars in the figure are 2 µm.

### The N-terminal region of ISAP1 is important for localization and function

Aside from its two CC domains (Fig. 4A, B), ISAP1 lacks homology to known proteins or domains that would provide a clue towards its function. We have previously shown the importance of CC domains in the early daughter protein IMC32 and the IMC cytoskeletal protein ILP1 (23, 24). To determine if the ISAP1 CC domains are important for targeting or function, we deleted each of the domains individually or together and expressed the deletion constructs in Δ*isap1* parasites (Fig. 4A). All of the deletions localized to a series of faint spots similar to that seen for the endogenously tagged protein (Fig. 4C). Surprisingly, all of the deletion constructs were also fully able to rescue the knockout as assessed by plaque assays (Fig. 4D). These results demonstrate that the CC domains of ISAP1 are dispensable for its localization and function.

**Figure 4.**
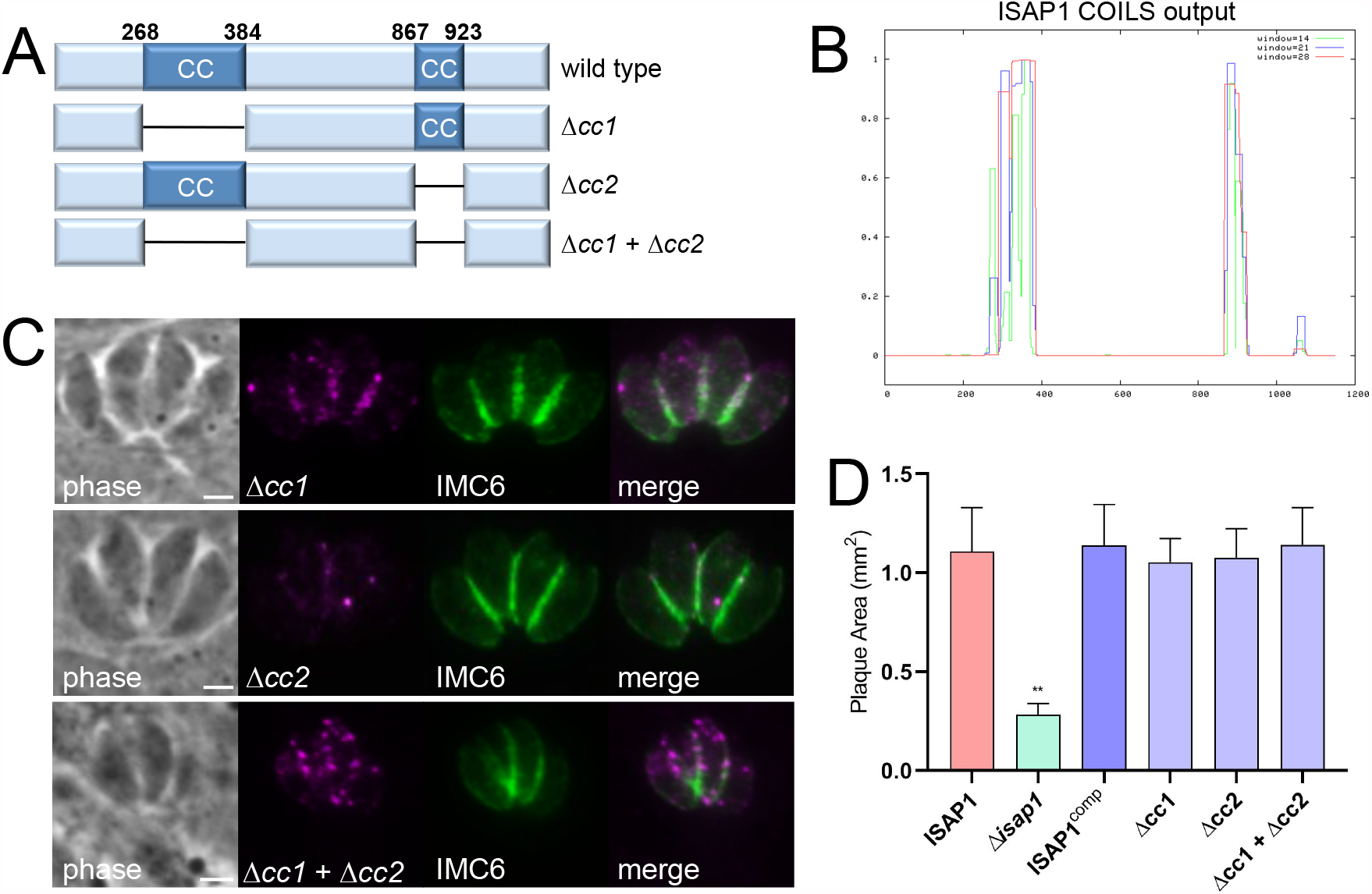
The CC domains of ISAP1 are not required for function. A) Diagram showing deletions of the CC domains individually or together. B) ISAP1 CC domains predicted by the COILS Server (43). C) IFA showing that the CC deletions localize to punctae similar to wild-type ISAP1. Magenta: mouse anti-HA; Green: rabbit anti-IMC6. Scale bar = 2 µm. D) Plaque assays showing that ISAP1 CC deletion constructs can fully rescue the plaque defect of Δ*isap1* parasites (**, *P* < 0.01).

We then focused on the N-terminal region upstream of the first CC domain (residues 1-250) and the C-terminal region downstream of the second CC domain (residues 1041-1148), as these portions have higher regions of homology (Fig. S1), suggesting that they may contain functional domains (Fig. 5A). While the C-terminal deletion construct rescued the knockout, absence of the N-terminal region (Δ2-250) eliminated the ability to rescue function. To further dissect the functional region of the N-terminal domain, we made two additional smaller deletions that removed residues 2-121 or 2-180, each of which result in the loss of regions with significant homology (Fig. S1). Neither of these constructs were able to rescue the phenotype of the knockout, demonstrating that the N-terminal 121 amino acids are essential for ISAP1 function (Fig. 5B). To determine if this was due to a loss of trafficking, we expressed the Δ2-121 and Δ2-250 ISAP1 constructs with a smMYC tag in wild-type parasites in which endogenous ISAP1 was tagged with smHA. IFA analysis showed that the deletion constructs failed to localize to the punctae, but instead localized primarily to the periphery of the parasite, consistent with IMC localization (Fig. 5C). Detergent extraction experiments showed that these deletion proteins still associated with the cytoskeleton, demonstrating that these regions are not necessary for cytoskeletal tethering (Fig. 5D). Together, this data demonstrates that the N-terminal 121 amino acids of ISAP1 are required for sutures punctae localization and function, but not for association with the IMC cytoskeleton.

**Figure 5.**
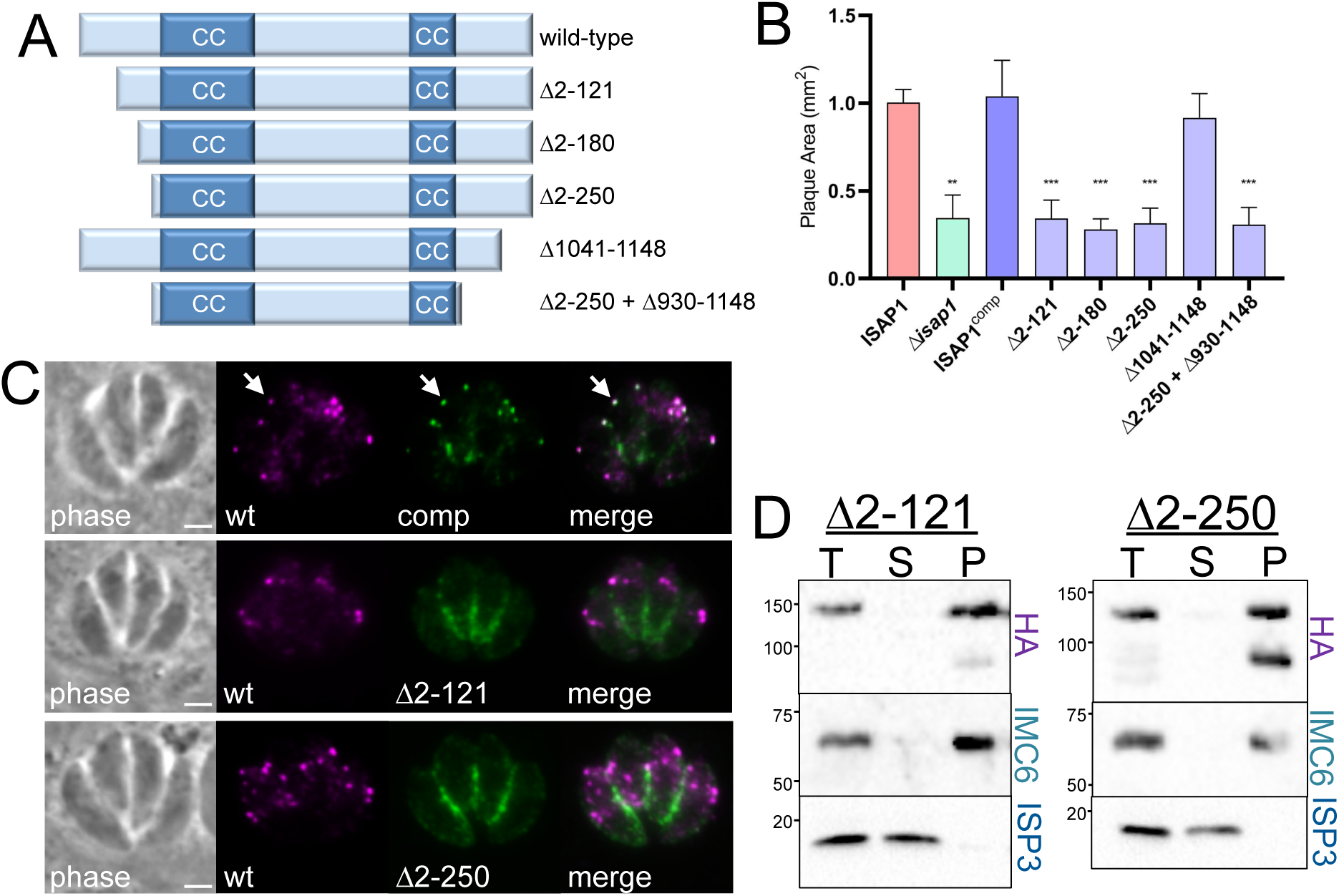
The N-terminal region of ISAP1 is important for punctae trafficking and function. A) Diagram showing the N and C-terminal deletion constructs. B) Plaque assays showing that the conserved C-terminal region of ISAP1 can be deleted but the N-terminal 121 amino acids of the protein is important for function (**, *P* < 0.01; ***, *P* < 0.001). C) IFA showing the full length smMyc-tagged ISAP1 complement (comp) targets to punctae similar to the endogenously smHA-tagged protein (wt) (arrows). Loss of the N-terminal 121 or 250 amino acids of ISAP1 results in relocalization to the periphery of the parasite, consistent with IMC localization. Magenta: rabbit anti-HA; Green: mouse anti-Myc. Scale bar = 2 µm. D) Detergent fractionation showing that Δ2-121 and Δ2-250 constructs still associate with the cytoskeleton. ISP3 and IMC6 are used as controls for the membrane and cytoskeletal fractions, respectively.

### *In vivo* biotinylation reveals candidate ISAP1-interacting proteins

To better understand how ISAP1 is tethered to the parasite’s cytoskeleton and functions in invasion and replication, we carried out *in vivo* biotinylation (using BioID2) with ISAP1 as a bait protein (Fig. 6A) (25). The endogenously tagged ISAP1-BioID2 fusion protein localized to spots in the cytoplasm similar to the HA-tagged protein (Fig. 6B). The fusion was also active as assessed by streptavidin staining upon the addition of biotin to the media, although the staining was low in agreement with the weak detection of the endogenously tagged bait protein. To identify candidate interacting proteins, we performed a large scale ISAP1-BioID2 *in vivo* biotinylation experiment. We have previously shown with the cytoskeletal IMC suture protein ISC4 that detergent fractionation can dramatically reduce background (13), thus we included this step for the ISAP1-BioID2 experiment and analyzed the purified proteins via mass spectrometry.

**Figure 6.**
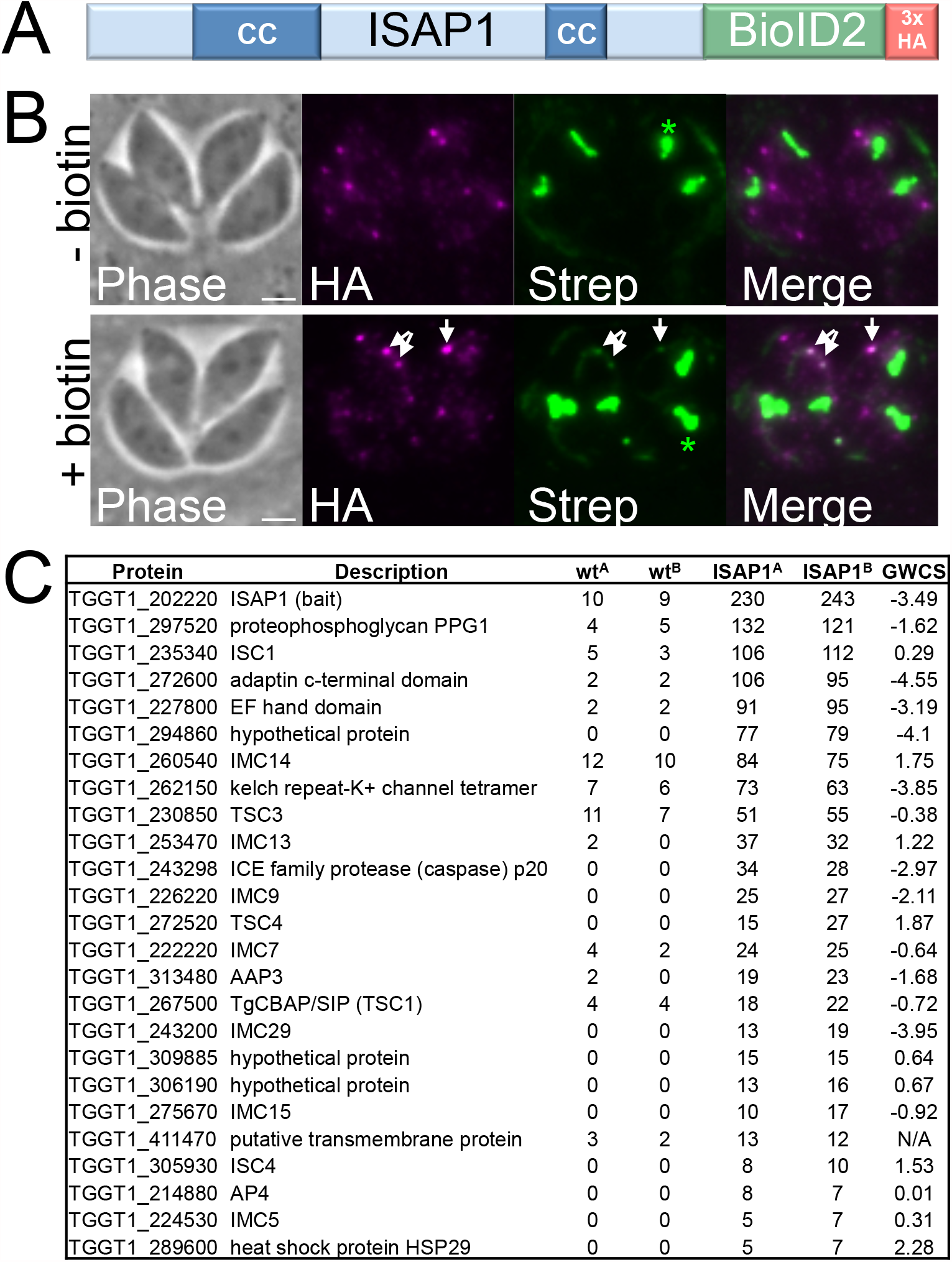
*In vivo* biotinylation using ISAP1 as bait identifies candidate interacting proteins. A) Diagram showing the ISAP1-BioID2 fusion generated by endogenous gene tagging. A 3xHA tag is included for detection of the fusion protein. B) IFA showing that the ISAP1-BioID2 fusion targets to punctae similar to the wild-type protein and is active as assessed by faint streptavidin staining upon addition of biotin to the media (arrows). Asterisks note the endogenously biotinylated signal in the apicoplast. Magenta: mouse anti-HA; Green: streptavidin 488. Scale bar = 2 µm. C) Table showing the top 25 hits from streptavidin purification of ISAP1-BioID2 following detergent fractionation of the cytoskeleton. Two replicates were performed (A, B samples) and untagged parasites plus biotin were used as the control.

Confirming the activity of the bait protein, ISAP1 was the top hit identified by mass spectrometry (Fig. 6C). In addition, 14 of the top 25 hits were known IMC proteins including IMCs 4/5/7/9/13/14/15/29, the IMC suture proteins ISC1/ISC4/TSC1 (CBAP/SIP)/TSC3/TSC4, and the apical annuli protein AAP3. Also highly ranked in this dataset were TgGT1_297520 which is annotated as proteophosphoglycan 1 (PPG1), and two proteins which have been implicated in vesicle transport, the intersectin-1-like protein TgGT1_227800 and the putative AP-2 adaptor complex member TgGT1_272600 (26). TgGT1_297520 (PPG1) was previously reported to be undetectable in tachyzoites but localizes to cytoplasmic spots plus the parasitophorous vacuole in bradyzoites (27). The localization of TgGT1_227800 has not been shown, but TgGT1_272600 reportedly localizes to low abundance cytoplasmic puncta that colocalize with the vesicle trafficking protein DrpC (although this data was not shown) (26, 27). These results together suggest potential links between ISAP1 with the IMC cytoskeleton and identifies several candidate interactors in the IMC punctae.

### Verification of BioID hits identifies proteins that colocalize with ISAP1

To determine if TgGT1_297520 (PPG1), TgGT1_227800 (intersectin-1-like), and TgGT1_272600 (AP-2) represent likely ISAP1 partners, we used CRISPR/Cas9 to endogenously tag their genes in a spaghetti monster ISAP1-tagged background (22, 28). IFA analysis showed that TgGT1_227800, TgGT1_297520, and TgGT1_272600 all localized in cytoplasmic spots that colocalized well with ISAP1 (Fig. 7A-C). To determine if these proteins are associated with the cytoskeleton, we conducted detergent extraction experiments as done above for ISAP1. While we were unable to detect TgGT1_272600 in the diluted conditions required for fractionation, both TgGT1_297520 and TgGT1_227800 fractionated primarily with the cytoskeletal fraction, although both proteins reproducibly suffered significant breakdown during fractionation (Fig. 7D, E). The colocalization and cytoskeletal cofractionation of these players agrees with their high ranking in the BioID experiment and suggests that they may interact with ISAP1 at this location.

**Figure 7.**
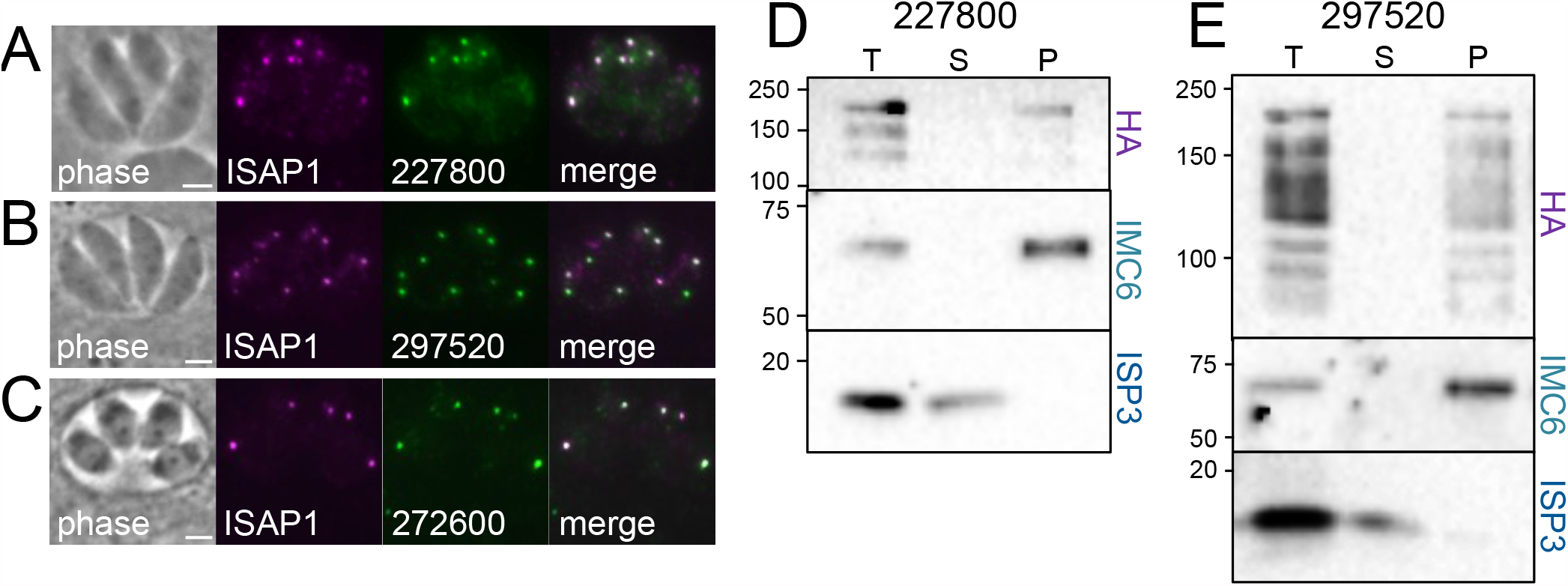
Endogenous tagging of candidates reveals proteins that colocalize with ISAP1. A) IFA showing the intersectin-1-like protein TgGT1_227800-smOLLAS localizes to discrete spots that colocalize with ISAP1-smHA. Magenta: rabbit anti-HA. Green: rat anti-OLLAS. Scale bar = 2 µm. B) IFA showing that TgGT1_297520-3xHA also colocalizes with the ISAP1-smOLLAS punctae. Magenta: rat anti-OLLAS; Green: mouse anti-HA. Scale bar = 2 µm. C) The AP2 adaptor complex member TgGT1_272600-3xHA also colocalizes with ISAP1-smOLLAS. Magenta: rat anti-OLLAS; Green: mouse anti-HA. Scale bar = 2 µm. D, E) Detergent fractionation shows that TgGT1_227800 and TgGT1_297520 are tethered to the cytoskeleton, although both of the proteins are labile and suffered substantial reproducible breakdown during fractionation. ISP3 and IMC6 are used as controls as above.

### The vesicle trafficking protein DrpC partially colocalizes with ISAP1 and largely fractionates with the cytoskeleton

TgGT1_227800 and TgGT1_272600 were previously identified as co-precipitating proteins with the vesicle trafficking protein DrpC, which also localizes to a series of cytoplasmic spots in the parasite (26). While DrpC was not identified in our BioID experiments (and ISAP1 was not identified in the DrpC pulldown), its similar localization pattern and common partners suggested that it may also colocalize with ISAP1. We thus assessed DrpC colocalization in the ISAP1-smOLLAS tagged strain. Most of the DrpC punctae colocalize with ISAP1, but additional spots were also observed (Fig. 8A). Detergent fractionation also demonstrated that a substantial portion of DrpC partitions with the cytoskeleton, although some is released by detergent extraction (Fig. 8B). We were surprised that DrpC and other putative trafficking proteins were tethered to the cytoskeleton, thus we also examined the related protein DrpB (29), and found it was efficiently released into the detergent soluble fraction, as expected (Fig. 8C). Together this data further links vesicle trafficking elements to ISAP1 in the parasite’s cytoskeleton and suggests that the sutures punctae are a site of trafficking of cargo across the IMC.

**Figure 8.**
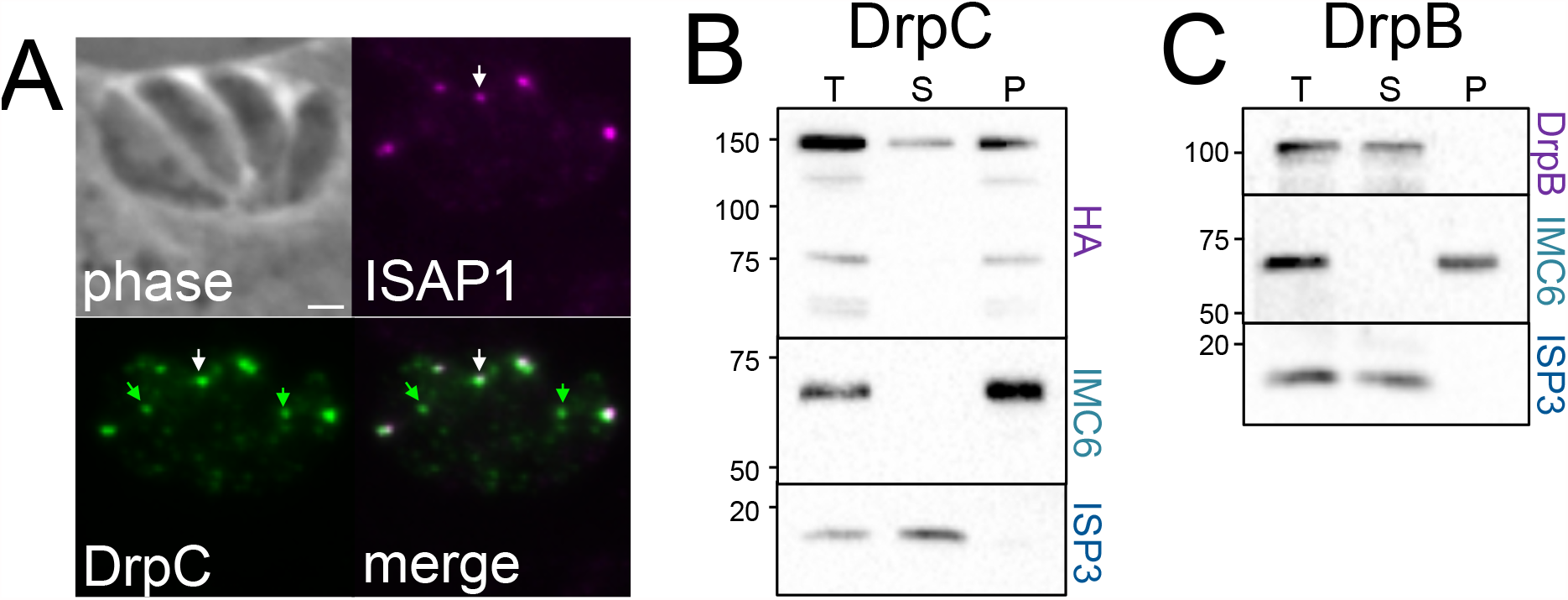
The membrane trafficking protein DrpC colocalizes with ISAP1 and associates with the cytoskeleton. A) IFA showing that DrpC-3xHA mostly colocalizes with ISAP1-smOLLAS (white arrows), though unique DrpC spots are also present (green arrows). Magenta: rat anti-OLLAS; Green: rabbit anti-HA. Scale bar = 2 µm. B) Detergent fractionation shows that a substantial portion of DrpC is surprisingly tethered to the cytoskeleton. ISP3 and IMC6 are used as controls as above. C) The related dynamin-related protein DrpB is readily solubilized by detergent extraction.

## Discussion

In this paper, we identify and characterize ISAP1, a new component of the *T. gondii* IMC that localizes to distinct cytoskeletal punctae that are embedded in the sutures of the organelle. Low expression levels made our initial characterization of the protein difficult to interpret, but this was resolved using high sensitivity spaghetti-monster tags that enabled better detection of the protein (20, 28). The increase in ISAP1 expression during daughter bud formation suggests that the punctae are synthesized and assembled onto the forming daughter buds with a “just in time” approach as is seen for other IMC proteins (11). This spot-like pattern anchored in the sutures thus represents a new type of localization within the IMC.

Like most IMC proteins, ISAP1 lacks homology to known proteins or identifiable domains other than its CC domains. CCs are structural motifs that consist of alpha helices that are coiled together and often function in oligomerization or protein-protein interactions (18). We have previously shown that CCs play important roles in the IMC for the cytoskeletal protein ILP1 and the early daughter membrane protein IMC32, and these motifs are also frequently present in the constituents of the conoid and apical annuli in *T. gondii* (15, 17, 23, 24). Surprisingly, deletion of the ISAP1 CCs individually or together demonstrated that they are dispensable for localization and function. However, our deletion analyses revealed a conserved domain in the N-terminal 121 amino acids that plays a critical role in both localization and function. While loss of this region disrupts punctae localization, the deletion protein retains tethering to the IMC cytoskeleton. Together, our deletion analyses suggest that IMC cytoskeletal binding is likely conferred by the central region of the protein between the CC domains (or by residues 923-1041 downstream of the second CC), with the N-terminal region responsible for tethering to the punctae, perhaps via binding to another protein at this location.

Our BioID experiments revealed a number of candidate interactors that may mediate binding to the IMC cytoskeleton or punctae organization. The IMC suture proteins ISC1/ISC4/TSC1/TSC3/TSC4 were highly ranked in the BioID experiment, which may represent cytoskeletal attachment points on the longitudinal or transverse sutures (12). The array of known IMC proteins identified (e.g. IMC 7/9/13/14) may represent additional contact points that tether ISAP1 to the cytoskeleton. The small plaque size and replication defects of Δ*isap1* parasites are reminiscent of the knockout of the suture protein ISC3, although ISC3 is membrane associated and not cytoskeletal like ISAP1 (13). While ISC3 localization does not appear to be impacted in Δ*isap1* parasites, the loss of the cytoskeletal ISCs 1/2/4 may impact the ability of ISC3 or other membrane suture proteins to be properly tethered to the cytoskeleton (12, 13, 22). Loss of ISAP1 also appears to impact the structure or integrity of the cytoskeleton, which likely results in the morphological changes leading to replication and invasion defects. The precise role of how ISAP1 controls parasite shape is likely to be best understood by determining how the protein acts with its interactors within the sutures punctae.

One of the top BioID hits that colocalizes with ISAP1 and fractionates with the cytoskeleton was TgGT1_297520, which is annotated the putative phosphoproteoglycan PPG1 (27). However, its similarity to phosphoproteoglycans is not clearly supported by BLAST or the OrthoMCL database (19, 30). Intriguingly, the protein contains a C-terminal GAR domain, which is involved in microtubule binding and could confer binding to the subpellicular microtubules or perhaps to the alveolins that form the IMC cytoskeleton (31). In addition, the protein structure-based prediction program iTasser suggests similarity to the cytoskeletal protein talin, which is known to interact with vimentin, an intermediate filament protein like the alveolins (32, 33). Also identified were the membrane trafficking implicated proteins TgGT1_227800 (intersectin-1-like) and TgGT1_272600 (AP-2 complex member) that were previously identified in DrpC pulldowns (26). The association with membrane trafficking proteins suggests that they may function in the delivery of cargo into or across the IMC. The association with the AP-2 adaptor complex, intersectin-1-like proteins, and DrpC could also suggest a role in endocytosis at this site. Endocytosis is still poorly understood in *T. gondii*, but it is believed to occur at a structure called the micropore (26, 34). The IMC sutures punctae seem likely to correlate with the micropore, but additional experiments will be necessary to confirm this and a direct role in endocytosis at this site.

While DrpC colocalizes partially with ISAP1 in the IMC sutures punctae, it was not identified in our ISAP1-BioID2 experiment. This may indicate that it is more distantly located in the punctae or biotinylation was impeded by the other colocalizing proteins which may serve as a bridge between ISAP1 and DrpC. Because we used detergent fractionation to enrich for cytoskeletal factors, it is also possible that other interacting or proximal proteins were missed due to the fractionation, which would best be evaluated using whole parasite extracts and the improved TurboID (35). Overall, our data agrees perfectly with a recent study that demonstrated that DrpC localizes in a spot-like pattern like ISAP1, associates with membrane trafficking proteins, and regulates IMC structure (26). Together, these studies may indicate that these proteins serve as a portal for trafficking of material across the IMC at discrete points along the IMC sutures. DrpC has additionally been shown to function in mitochondrial fission at the end of replication (36). It is possible that DrpC is able to perform multiple functions such as mitochondrial fission via its localization to additional punctae within the parasite. Further dissection of ISAP1, DrpC and each of their interacting partners will provide a deeper understanding into how these proteins regulate cellular functions in *T. gondii*.

## Materials and Methods

### *Toxoplasma* and host cell culture

*T. gondii* RHΔ*ku80*Δ*hpt* and modified strains were grown on confluent monolayers of human foreskin fibroblast (HFF) host cells in DMEM supplemented with 10% fetal bovine serum, as previously described (37).

### Antibodies

The following previously described primary antibodies were used in immunofluorescence (IFA) or Western blot assays: mouse anti-ISP1 (38), mouse anti-ISP3 (39), rabbit anti-IMC6 (23), anti-F1β subunit (mAb 5F4) (40), and anti-ATrx1 (mAb 11G8) (41). The hemagglutinin (HA) epitope was detected with mouse anti-HA (mAb HA.11) (BioLegend) or rabbit anti-HA (Invitrogen). The c-Myc epitope was detected with mouse anti-Myc (mAb 9E10) and the OLLAS tag was detected using the rat monoclonal anti-OLLAS (28). For production of ISC1 and ISC2 antibodies, the complete coding sequences of the genes were cloned into the pET28 and pET160 bacterial expression vectors, respectively. The constructs were transformed into BL21(DE3) *E. coli*, and proteins were induced with 1 mM IPTG and purified using Ni-NTA agarose under denaturing conditions as described (42). The samples were then dialyzed into PBS to remove the urea, and rat antisera against the proteins were produced by Cocalico Biologicals.

### Immunofluorescence assays (IFA) and Western blot

For IFA, HFFs were grown to confluency on coverslips and infected with *T. gondii* parasites. After 18-36 hours, the coverslips were fixed and processed for indirect immunofluorescence using either 3.7% formaldehyde or 100% ice-cold methanol as previously described (42). Primary antibodies were detected by species-specific secondary antibodies conjugated to Alexa 594/488. The coverslips were mounted in Vectashield (Vector Labs) and viewed with an Axio Imager.Z1 fluorescence microscope (Zeiss) as described (43).

For Western blot, parasites were lysed in Laemmli sample buffer (50 mM Tris-HCl [pH 6.8], 10% glycerol, 2% SDS, 0.1M DTT, 0.1% bromophenol blue) and lysates were resolved by SDS-PAGE and transferred onto nitrocellulose membranes. Blots were probed with the indicated primary antibodies, followed by secondary antibodies conjugated to horse radish peroxidase (HRP). Target proteins were visualized by chemiluminescence (Thermo Scientific).

### Epitope tagging

For endogenous tagging of proteins, we used CRISPR/Cas9 as previously described (16, 22). The appropriate guides were ligated into the pU6 Universal plasmid and the tag plus selectable marker was amplified from LIC tagging plasmids with 40bp flanking regions for recombination at the 3’ end of each gene. Following transfection of the guide plus PCR product, transgenic parasites were selected in the appropriate drug media (containing either 1 µM pyrimethamine, 50 µg/ml mycophenolic acid/xanthine, or 1 µM chloramphenicol) and cloned by limiting dilution. Clones that had undergone the intended recombination event were screened by IFA and Western blot against the epitope tag. Tagging of TgGT1_248740, ISC5, and TSCs 2-6 was performed using the LIC method as previously described (13). The PCR verification of ISC4 tagging in the Δ*isap1* parasites was performed with primers p64-p66 (Table S1).

### Detergent extractions

Extracellular parasites were washed in PBS, pelleted, and lysed in 1 mL of 1% Triton X-100 lysis buffer (50mM Tris-HCl [pH 7.4], 150mM NaCl) supplemented with Complete Protease Inhibitor Cocktail (Roche) for 20 min on ice. Lysates were centrifuged for 10 min at 16,000 x *g* at 4°C. Equivalent amounts of total, supernatant (detergent soluble), and pellet (detergent insoluble) fractions were separated by SDS-PAGE and analyzed by Western blot. IMC6 and ISP3 served as controls for the cytoskeletal and membrane fractions, respectively (23, 39).

### Gene knockout and phenotypic analyses

CRISPR/Cas9 and homologous recombination was used to knockout the *ISAP1* gene, as previously described using primers p5-p10 (44). Knockout clones were verified by IFA and PCR using primers p11-p14.

For plaque assays, intracellular parasites were collected by scraping and passaging through a 27-gauge needle, and then equivalent parasite numbers were allowed to infect HFF monolayers. 7 days after infection, cells were fixed with 100% ice-cold methanol and stained with crystal violet (44). The area of 50 plaques per condition was measured using ZEN software (Zeiss). All plaque assays were performed in triplicate using biological replicates. Plaque number was counted manually to measure efficiency of plaquing. Statistical significance was calculated using unpaired t-tests comparing each condition against the wild type HA-tagged ISAP1.

For egress assays, parasites were grown on coverslips with HFF monolayers for 30 hours, washed with warm HBSS, incubated with A23187 or DMSO control at 37°C for 3 min, and then fixed and stained, as previously described (9). The percentage of lysed vacuoles was counted for three replicates per condition.

For invasion assays, parasites were settled onto HFF monolayers on coverslips in warm Endo buffer for 20 min, and then allowed to invade using warm D1 media for 15 min, as previously described (9). Coverslips were fixed and blocked, and extracellular parasites were differentially stained before the cells were permeabilized and all parasites were stained, to record parasites as invaded or not. Invasion assays were performed in triplicates, and statistical significance was calculated using unpaired t-tests comparing each condition against the wild type HA-tagged ISAP1.

### Complementation with wild type and mutant constructs

To generate the wild type complement construct, the entire coding region of the gene was PCR amplified from genomic DNA and cloned into a UPRT-locus knockout vector driven by the ISC6 promoter, as previously described (primers p15-p18) (9). The phosphomutant was constructed using synthetic genes (Quintara Biosciences) to mutate each phosphosite into an alanine in the complementation vector. Deletion constructs were built with the complementation vector as a template using the Q5 Site-Directed Mutagenesis Kit to delete specific sequences (primers p19-p31). The plasmids were linearized and transfected into the Δ*isap1* parasites and selected with 5-Fluoro-5’-deoxyuridine (FUDR). Parasites expressing the complementation constructs were cloned by limiting dilution.

### Affinity purification of biotinylated proteins

HFF monolayers infected with parasites expressing the ISAP1-BioID2 fusion or the respective parental line were grown in media containing 150 µM biotin for 24 hours prior to parasite egress. Approximately 10^9^ extracellular parasites were collected, washed in PBS, and lysed in RIPA buffer (50 mM Tris [pH 7.5], 150 mM NaCl, 0.1% SDS, 0.5% sodium deoxycholate, 1% NP-40) supplemented with Complete Protease Inhibitor Cocktail (Roche) for 30 minutes on ice. Lysates were centrifuged for 15 min at 16,000 x *g* to pellet insoluble debris and then incubated with High Capacity Streptavidin Agarose (Pierce) at room temperature for 4 hours under gentle agitation. Beads were collected by centrifugation and washed five times in RIPA buffer, followed by three washes in 8M urea buffer (8M Urea, 50mM Tris-HCl [pH 7.4], 150mM NaCl). 10% of each sample was boiled in Laemmli sample buffer and eluted proteins were analyzed by Western blot while the remaining material was digested directly from the beads for mass spectrometry as described (12).

### Biotinylated protein sample digestion and desalting

The proteins bound to streptavidin beads were reduced and alkylated via sequential 20-minute incubations of 5mM TCEP and 10mM iodoacetamide at room temperature in the dark while being mixed at 1200 rpm in an Eppendorf thermomixer. Proteins were then digested by the addition of 0.1µg Lys-C (FUJIFILM Wako Pure Chemical Corporation, 125-05061) and 0.8µg Trypsin (Thermo Scientific, 90057) while shaking at 37°C overnight. The digestions were quenched via addition of formic acid to a final concentration of 5% by volume. Each sample was desalted via C18 tips (Thermo Scientific, 87784) and then resuspended in 15µL of 5% formic acid before analysis by LC-MS/MS.

### LC-MS Acquisition and Analysis

Peptide samples were separated on a 75uM ID, 25cm C18 column packed with 1.9µM C18 particles (Dr. Maisch GmbH) using a 140-minute gradient of increasing acetonitrile and eluted directly into a Thermo Orbitrap Fusion Lumos instrument where MS/MS spectra were acquired by Data Dependent Acquisition (DDA). Data analysis was performed using the ProLuCID (45) and DTASelect2 (46, 47) algorithms as implemented in the Integrated Proteomics Pipeline - IP2 (Integrated Proteomics Applications, Inc., San Diego, CA). Protein and peptide identifications were filtered using DTASelect and required a minimum of two unique peptides per protein and a peptide-level false positive rate of less than 1% as estimated by a decoy database strategy.

## Acknowledgements

We thank Silvia Moreno for sharing the HA and Myc tag spaghetti monster epitope tagging constructs and Gary Ward for providing IMC1 antibodies. We also thank members of the Bradley lab for reading and editing of the manuscript. This work was supported by the Undergraduate Research Scholars Program (URSP), the Undergraduate Research Fellows Program (URFP) and Whitcome fellowships to J.H.C, as well as NIH grants #AI064616 to P.J.B. and #GM089778 to J.A.W. The funders had no role in the study design, data collection, interpretation, or decision to submit the work for publication.

## Supplemental Figure Legends

**Fig. S1.**
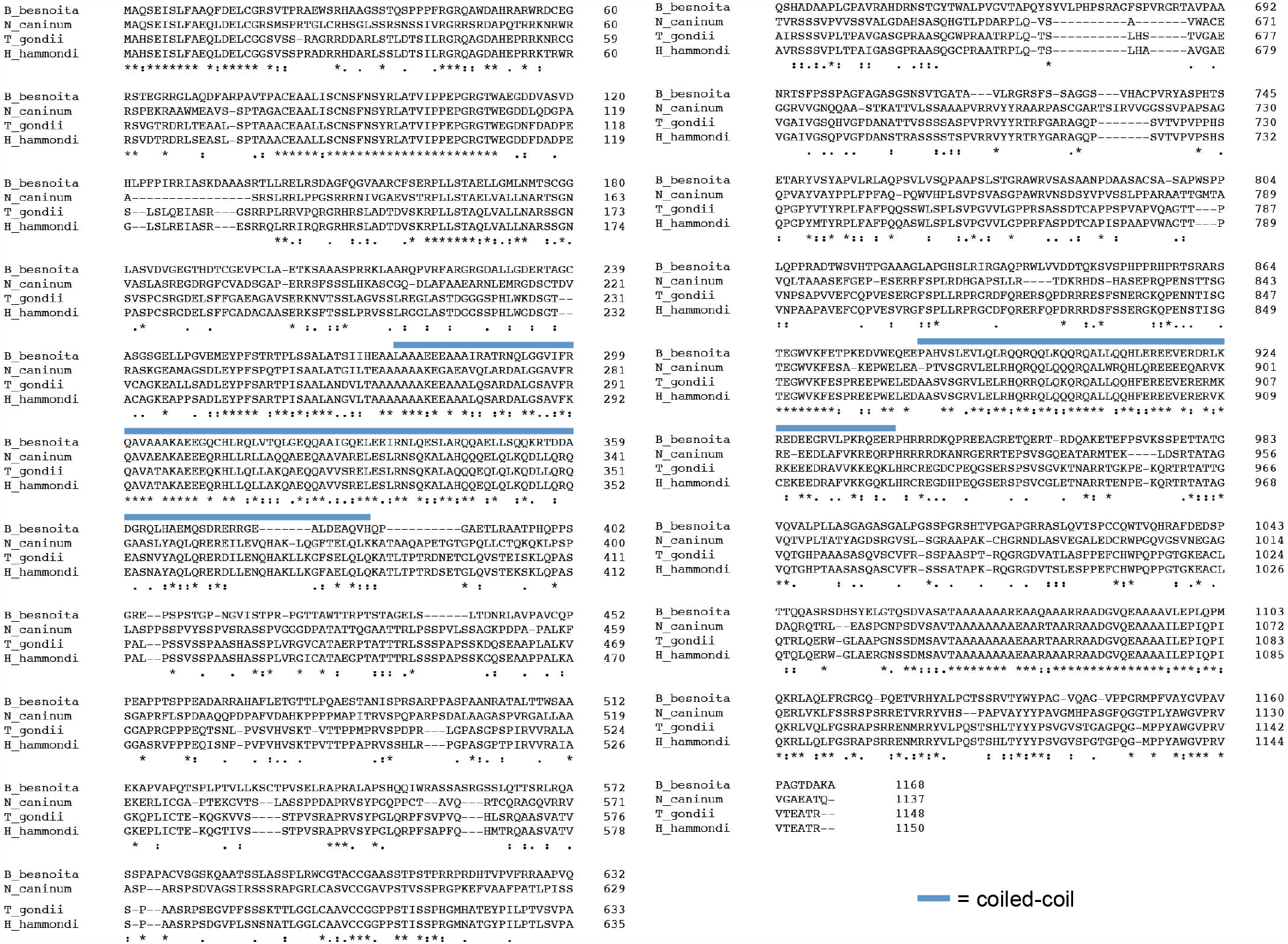
Alignment of TgGT1_202220 orthologues. Clustal Omega alignment of TgGT1_202220 orthologues from *H. hammondi, N. caninum*, and *B. besnoita* (19, 48). An asterisk indicates a fully conserved residue, whereas a colon indicates strong conservation and a period indicates weaker similarity. The positions of the coiled-coil domains are noted above the alignment with thick blue lines.

**Fig. S2.**
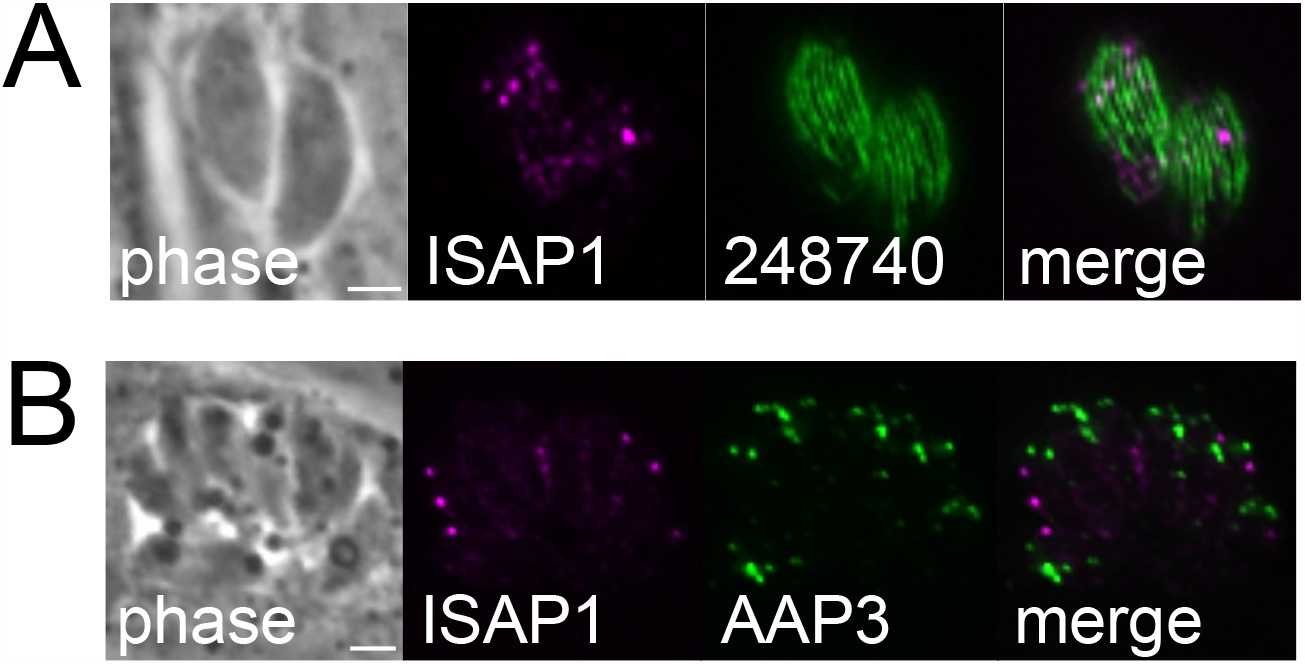
ISAP1 does not colocalize with the subpellicular microtubules or apical annuli. A) IFA showing that ISAP1-smOLLAS does not colocalize with the subpellicular microtubules seen by HA-tagging TgGT1_248740, which localizes to the subpellicular microtubules beneath the apical cap (13). Magenta: rat anti-OLLAS; Green: rabbit anti-HA. Scale bar = 2 µm. B) IFA showing that ISAP1-smOLLAS does not colocalize with the apical annuli observed via AAP3 HA-tagging (15). Magenta: rat anti-OLLAS; Green: rabbit anti-HA. Scale bar = 2 µm.

**Fig. S3.**
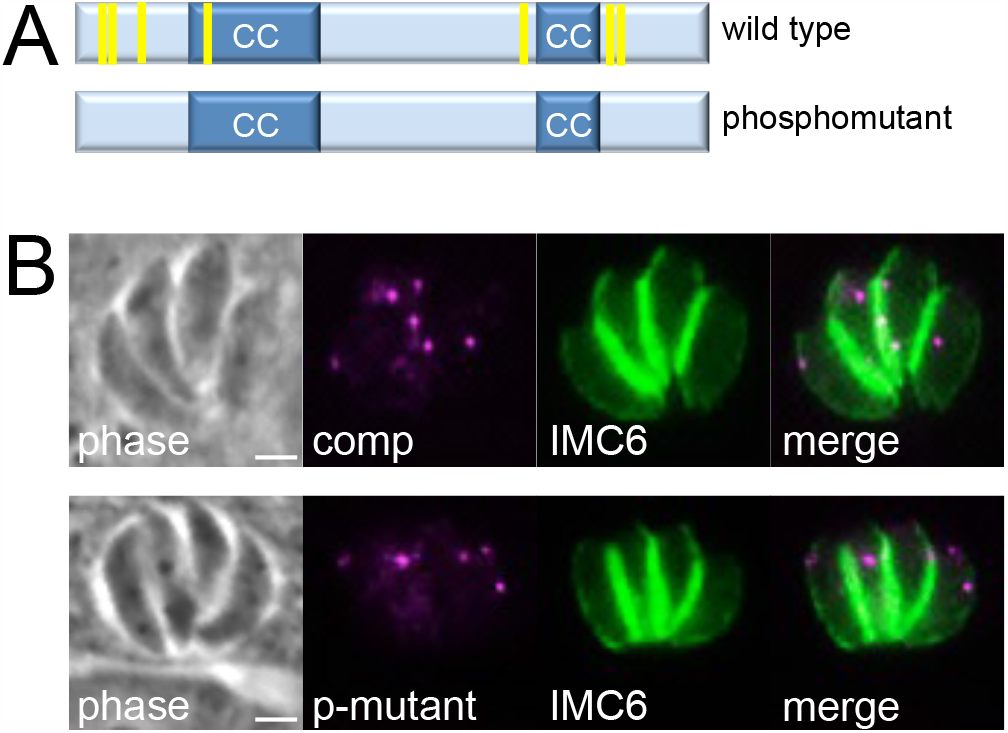
Mutagenesis of the ISAP1 phosphorylation sites. A) Diagram showing the position of ISAP1 phosphorylation sites (yellow bars, residues S74, T76, T149, S280, S856, S936, S941) identified by phosphoproteomics. All 7 serine or threonine residues were mutated to alanine (19, 49). B) IFA showing that the ISAP1 phosphomutant localizes to cytoplasmic punctae similar to the wild-type complement protein. Rescue of the Δ*isap1* plaque defect by the phosphomutant is shown in Fig. 2E. Magenta: mouse anti-HA; Green: rabbit anti-IMC6. Scale bars = 2 µm.

## Supplemental Table Legends

**Table S1.**
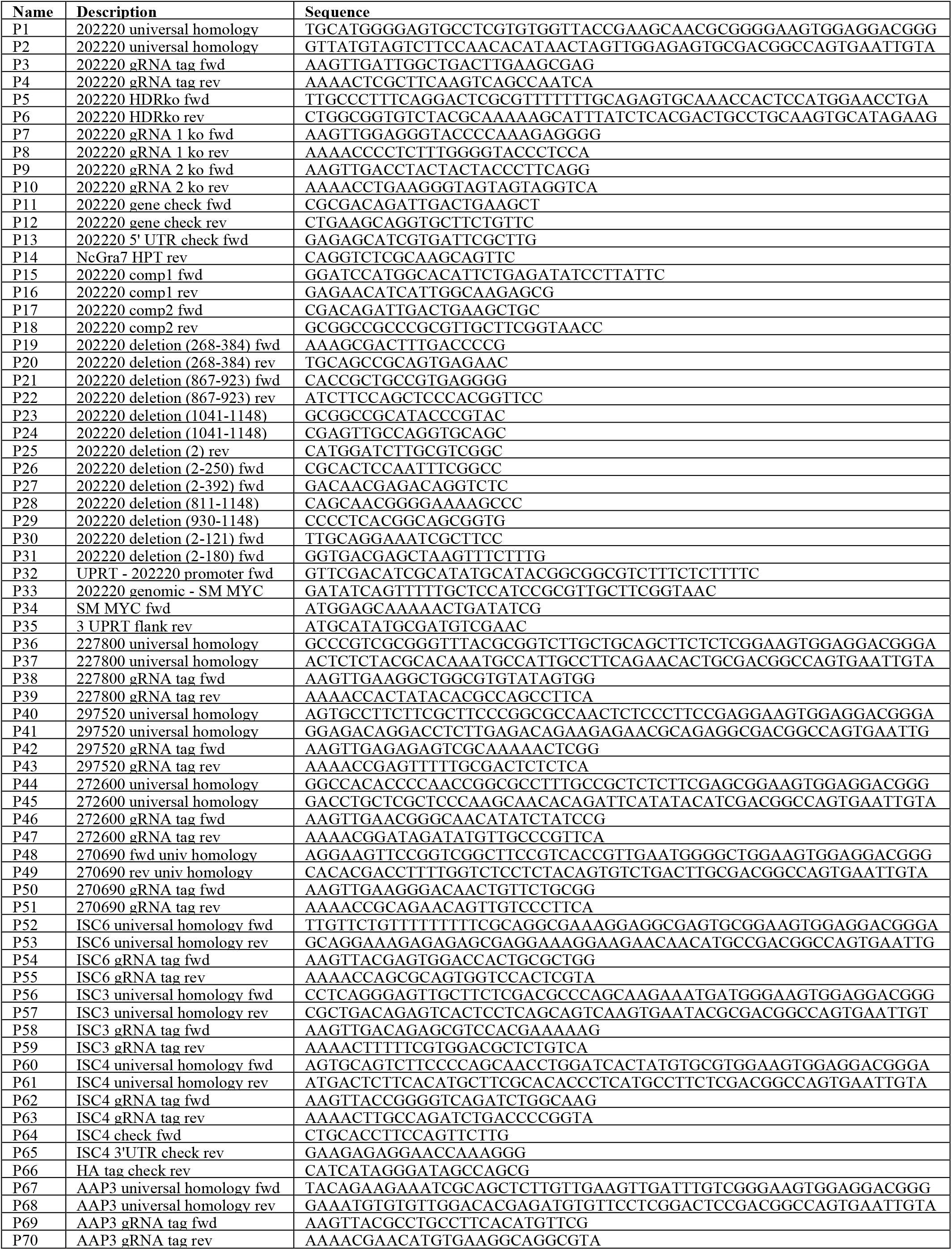
Oligonucleotide primers used in this study. All primer sequences are shown in the 5’ to 3’ orientation.

